# A novel gene-by-environment quantitative trait locus on mouse chromosome 15 underlies susceptibility to acute ozone-induced lung injury

**DOI:** 10.1101/2021.05.20.445039

**Authors:** Adelaide Tovar, Gregory J. Smith, Joseph M. Thomas, Kathryn M. McFadden, Samir N. P. Kelada

## Abstract

Respiratory toxicity caused by the common urban air pollutant ozone (O_3_) varies considerably within the human population and across inbred mouse strains, suggestive of gene-environment interactions (GxE). Though previous studies genetic mapping studies using classical inbred strains have identified several quantitative trait locus (QTL) and candidate genes underlying responses to O_3_ exposure, precise mechanisms of susceptibility remain incompletely described. We sought to expand our understanding of the genetic architecture of O_3_ responsiveness using the Collaborative Cross (CC) recombinant inbred mouse panel, which contains more genetic diversity than previous inbred strain panels. We evaluated hallmark O_3_-induced respiratory phenotypes in 56 CC strains after exposure to filtered air or 2 ppm O_3_, and performed focused genetic analysis of variation in lung injury as measured by the total bronchoalveolar lavage protein concentration. Because animals were exposed in sex- and batch-matched pairs, we defined a protein response phenotype as the difference in lavage protein between the O_3_- and FA-exposed animal within a pair. The protein response phenotype was heritable, and QTL mapping revealed two novel loci on Chromosomes 10 (peak: 26.2 Mb; 80% CI: 24.6-43.6 Mb) and 15 (peak: 47.1 Mb; 80% CI: 40.2-54.9 Mb), the latter surpassing the 95% significance threshold. At the Chr. 15 locus, C57BL/6J and CAST/EiJ founder haplotypes were associated with higher protein responses compared to all other CC founder strain haplotypes. Using additional statistical analysis and high-density SNP data, we delimited the Chr. 15 QTL to a ∼2 Mb region containing 21 genes (10 protein coding). Using a weight of evidence approach that incorporated candidate variant analysis, functional annotations, and publicly available lung gene expression data, we nominated three candidate genes (*Oxr1, Rspo2*, and *Angpt1*). In summary, we have shown that O_3_-induced lung injury is modulated by genetic variation and demonstrated the value of the CC for uncovering and dissecting gene-environment interactions.

## Introduction

Ozone (O_3_) is a potent oxidant gas and ground-level air pollutant. Acute O_3_ exposure causes temporary decrements in lung function (1, 2), respiratory inflammation (3, 4), and tissue injury (5, 6), as well as aggravating symptoms of common chronic lung diseases including asthma and chronic obstructive pulmonary disease (7–9). Short-term O_3_ exposure is also associated with an increased risk of respiratory tract infections and hospitalization (10). Importantly, these adverse outcomes have been linked with the pathogenesis of respiratory diseases (11). Thus, because of its involvement in disease incidence and exacerbations (12), identifying the molecular circuitry by which O_3_ exposure causes pulmonary inflammation and injury is a critical public health concern.

Controlled exposure and candidate gene studies in humans have provided strong evidence that responses to O_3_ vary widely and reproducibly across individuals, in a fashion partially attributable to genetic variation (13–19). Studies using panels of genetically diverse rodent strains have corroborated these findings, provided evidence of heritability, and identified multiple loci that control hallmark respiratory responses such as airway neutrophilia and injury (20, 21). These foundational studies proposed important candidate genes including *Tlr4, Nos2*, and *Tnf*, whose roles in O_3_ respiratory toxicity have since been thoroughly studied (22–27). Here, we sought to further characterize the genetic architecture of respiratory responses to acute O_3_ exposure, thereby illuminating novel gene-by-environment interactions (GxE) for future experimental validation and investigation.

For the current study, we used the Collaborative Cross (CC) genetic reference population, a multi-parental inbred strain panel derived by funnel inbreeding of five classical (A/J, C57BL/6J, 129S1SvImJ, NOD/ShiLtJ, NZO/H1LtJ) and three wild-derived mouse strains (CAST/EiJ, PWK/PhJ, WSB/EiJ) (28, 29). Nearly all previously published studies investigating genetic contributions to O_3_ responses focused on responses in classical laboratory strains, failing to capture the full breadth of genetic diversity available within the *Mus musculus* species. The CC captures over 90% of circulating genetic variants (> 40 million SNPs and several million indels and small structural variants), owing in large part to the inclusion of wild-derived strains from three different *Mus musculus* subspecies (*M. m. domesticus, castaneus* and *musculus*) (30). Moreover, the breeding strategy used to generate CC lines created novel allelic combinations, thus enhancing the range of phenotypic variation observed compared to that across the founder strains alone (31, 32). This population has already been used to identify genomic regions associated with various respiratory disease phenotypes including allergic inflammation in the airways (33–35) and susceptibility to virus-induced respiratory disease endpoints (36, 37).

Hence, this resource is well-suited for toxicogenomic discovery and identification of genes and variants contributing to O_3_-induced responses in the lung. We measured inflammatory and injury responses in mice from 56 CC strains after a 3-hour exposure to filtered air (FA) or 2 ppm O_3_. The analyses here are primarily focused on total protein concentration in the bronchoalveolar lavage fluid, a non-specific marker of lung injury, for which we identified two novel genomic loci (one significant and one suggestive) associated with variation in responsiveness. Using a weight of evidence approach along with available genome sequence data, we prioritized candidate genes within the significant locus, located on Chromosome 15.

## Materials and Methods

### Animals

Female and male mice from 56 Collaborative Cross strains were obtained from the UNC Systems Genetics Core Facility between August and November 2018. Mice were delivered as litter- and sex-matched pairs or trios (matches) upon weaning and aged on investigator’s racks until the time of exposure. Animals were housed in groups of three or more under normal 12-hour light/dark cycles, in polycarbonate cages with *ad libitum* food (Envigo 2929) and water on Teklad 7070C bedding (Envigo). All studies were conducted in compliance with a protocol reviewed by the University of North Carolina Institutional Animal Care and Use Committee in animal facilities approved and accredited by the Association for Assessment and Accreditation of Laboratory Animal Care International.

### Ozone exposure

All mice were 9-12 weeks of age at the time of filtered air or 2 ppm ozone exposure. Mice were randomly assigned to experimental groups within a pair or trio (referred to as a match), and all matches within a strain were randomized over the course of the study to address batch effects. A full calendar detailing the batches is included in Supplemental Figure S1. Mice were exposed to O_3_ in individual wire-mesh chambers without access to food or water, as described previously (38, 39). Exposures were performed at roughly the same time (from 9 am-12 pm), with highly stable concentrations of O_3_ delivered across all batches (Supplemental Figure S2).

### Phenotyping

#### Lung phenotyping

Mice were returned to their normal housing upon cessation of exposure for 21 hours, at which point mice were anesthetized via an i.p. injection of urethane (2 g/kg) and euthanized by exsanguination via the inferior vena cava/descending abdominal aorta. Two bronchoalveolar lavage (BAL) fractions were collected as described previously (39). Supernatant from the first BAL fraction was stored at −80°C for biochemical analysis. Pellets from both BAL fractions were pooled, and a portion was used for performing total and differential cell counts.

#### Total protein measurement

Total BAL protein was quantified using the Quant-iT Protein Assay kit (Thermo Scientific), using manufacturer’s instructions. Assays were performed in black 384-well plates using 1 μL of BAL fluid and 75 μL of assay reagent. Samples were randomly assigned to 1 of 5 plates and each sample was plated in triplicate. Fluorescence was measured using the Cytation 5 multi-mode plate reader (BioTek), and values were calculated from the mean of the triplicate compared to the standard curve.

### Statistical genetics analysis

#### Definition of ozone-induced lung injury phenotype using BAL protein data

Here, we define a protein response phenotype, indicative of lung injury, as the difference in total protein concentration between an O_3_-exposed mouse (or mice) and its FA-exposed match. For trios, the mean value from the two O_3_-exposed mice was used. Data from unpaired mice were excluded.

#### Heritability calculations

Broad-sense heritability (*H*^*2*^) was estimated using a Bayesian linear mixed model approach implemented in R/INLA (40), as described previously with Heterogeneous Stock rats (41). Importantly, because a strain-identity kinship matrix is used for the inbred CC strains to estimate genetic relatedness, additive and dominant genetic effects are confounded; thus, broad-sense rather than narrow-sense heritability is calculated, in contrast to the approach used in Keele *et al*. (41).

#### CC strain genotypes and haplotype mosaics

Inferred CC founder haplotype contributions were previously reconstructed by the UNC Systems Genetics Core using a Hidden Markov Model on weighted consensus genotype calls from the most recent common ancestors for a given CC strain. These founder genotype probability files are publicly available (http://csbio.unc.edu/CCstatus/index.py?run=FounderProbs). The genome cache was constructed with a reduced number of total loci, as described previously (42). Briefly, to minimize computational burden and reduce the testing penalty when performing quantitative trait locus (QTL) mapping, adjacent genomic regions with similar founder mosaics were identified and merged through averaging. The genome cache used for mapping was constructed with NCBI Build 37 positions, then converted to Build 38 positions using the UCSC Genome Browser liftOver function.

#### QTL mapping

Association between the protein response phenotype and the inferred CC founder haplotypes was performed at each point in the genome using a variant of the Haley-Knott regression termed regression on probabilities (ROP), as implemented in R/miqtl (42). First, a null model was built,

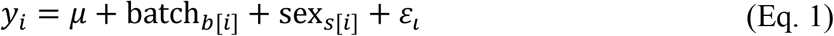

where *y*_*i*_ represents the phenotype value for an individual pair or trio, *i, μ* represents the population mean, variables for batch date (with 17 levels, *b* = 1…17) and sex of a given match are included as covariates. Then, a full model was built,

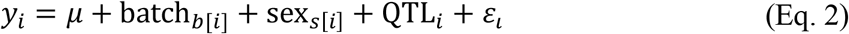

where QTL denotes the additive QTL effect of founder haplotypes at a given locus. The likelihoods of the two models were compared using a likelihood ratio test. QTL were deemed significant if they surpassed the genome-wide significance threshold, which was determined by permutation (n=1000).

#### QTL effect size and QTL location estimation

The variance attributed to a QTL was calculated by linear regression. Effect size was computed as

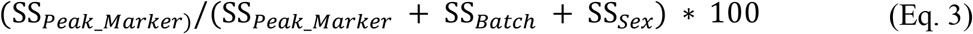

where SS is the sum of squares, or variation, attributed to founder haplotype at the peak marker (SS_*peak*___*Marker*_), batch (SS_*Batch*_), or sex (SS_*Sex*_). Confidence intervals for the QTL locations were defined using positional bootstrapping with R/miqtl. For each QTL, 1000 bootstraps were performed, and the 80% confidence interval was reported.

#### Haplotype effects estimation and allelic series inference

To estimate the effect of haplotype substitution at the detected QTL on protein response, we used Diploffect (43), a Bayesian linear mixed model that estimates confidence intervals for additive haplotype effects while incorporating uncertainty in haplotype contributions present in the genome cache used for mapping. To infer the allelic series at QTL, or number of functional alleles into which the founder haplotypes group, we used a Bayesian model selection method, Tree-based Inference of Multiallelism via Bayesian Regression (TIMBR) (44).

#### Candidate variant (merge) analysis

A multiallelic merge analysis was used to identify candidate variants within the 80% confidence interval of the significant QTL peak (45–47). This procedure tests whether a significant QTL signal, identified by the 8-haplotype association model, can be explained more parsimoniously by the pattern of known alleles at a given variant.

#### Weight of evidence criteria for candidate gene identification

To evaluate candidate genes within the QTL region, we used a weight of evidence approach with the following criteria:

1. Existence of variants (within or near a gene, identified by merge analysis) whose strain distribution pattern is similar to or the same as the haplotype effects from the eight-allele model used for mapping.
2. Presence of variants (ascertained from #1) that alter the coding portion of the transcript, with an emphasis on those that also alter or disrupt the amino acid sequence.
3. Evidence that the candidate gene is expressed in any compartment or cell type within the lung. For this criterion, we queried (1) our previously published bulk RNA-seq data (39) generated from conducting airways tissue and airway macrophages isolated from female C57BL/6J mice exposed to FA and 2 ppm O_3_, and (2) a variety of publicly available single-cell and bulk RNA-seq datasets using the EMBL-EBI Single Cell and Bulk Expression Atlases and LungMAP.
4. Relevant biological function, as judged by their entry in the Mouse Genome Informatics database and reference in previous publications.

#### Variant consequence prediction

SIFT (48) consequences were lifted from the Ensembl database, while PolyPhen-2 (49), PROVEAN (50) and SNAP2 (51) scores were calculated using default settings on their respective webtools.

### Data and code availability

Raw phenotypic data, metadata, and individual-level protein concentration data are available in Supplemental Table S1. All code used for described analyses is available in a single R file (‘tovar_2021.R’) in the Supplemental Material, except that used for merge analysis, which is available on GitHub (‘yanweicai/MergeAnalysisCC’). Relevant kinship matrices used for heritability calculations are provided in the Supplemental Material.

## Results

### Acute ozone (O_3_) exposure induces variable lung inflammation and injury across 56 Collaborative Cross (CC) strains

We utilized a design in which mice within a given Collaborative Cross (CC) strain were exposed to filtered air (FA) or ozone (O_3_) in sex- and litter-matched pairs or trios, hereafter referred to as matches (Figure 1). We observed an induction of neutrophilia that was highly variable across strains after O_3_ exposure (Figure 2). Using data from O_3_-exposed animals only (because most strains have no neutrophils after FA exposure), we estimated the broad-sense heritability (*H*^*2*^) to be 0.47 (95% CI: 0.36-0.60) and 0.52 (95% CI: 0.23-0.78) for percentage and total number of neutrophils in BAL, respectively. As a quantitative measure of lung injury, we measured total protein concentration in bronchoalveolar lavage (BAL) fluid (Figure 3A). This is a non-specific marker of serum contents (primarily serum albumin (4, 52)) that leak into the airspace upon damage to the alveolar epithelium and increased permeability of the underlying capillary beds. Evidence of modest strain effects were observed in FA-treated animals, while treatment effects were clear in O_3_-exposed animals. *H*^*2*^ was estimated at 0.26 (95% CI: 0.16-0.40) and 0.53 (95% CI: 0.41-0.64) for FA- and O_3_-exposed animals, respectively. We defined the total protein response as the difference in total protein concentration between the O_3_-exposed animal(s) and the FA-exposed animal for a given match within a strain. The range of total protein response values were nearly normally distributed, indicative of a polygenic, complex trait (Figure 3B), and *H*^*2*^ for this response trait was estimated at 0.39 (95% CI: 0.26-0.54).

**Figure 1.**
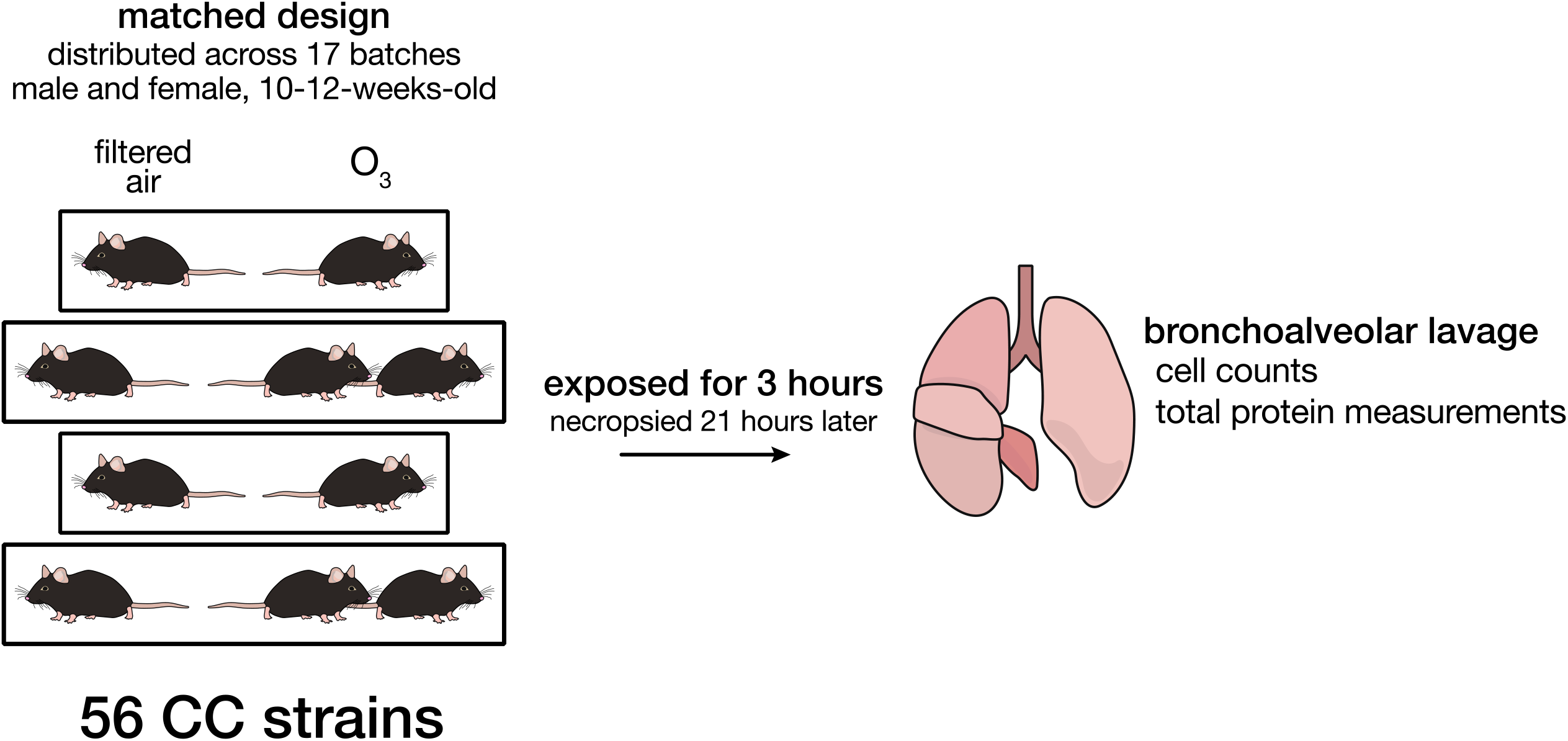
Study design. We exposed adult female and male mice from 56 Collaborative Cross (CC) strains in litter- and sex-matched pairs/trios to filtered air (FA) or 2 ppm ozone (O_3_) for 3 hours. Animals were necropsied 21 hours after exposure, and various parameters were measured including the type and quantity of inflammatory cells and the total protein concentration (lung injury marker) in the bronchoalveolar lavage (BAL) fluid. Matches were randomized over 17 batches.

**Figure 2.**
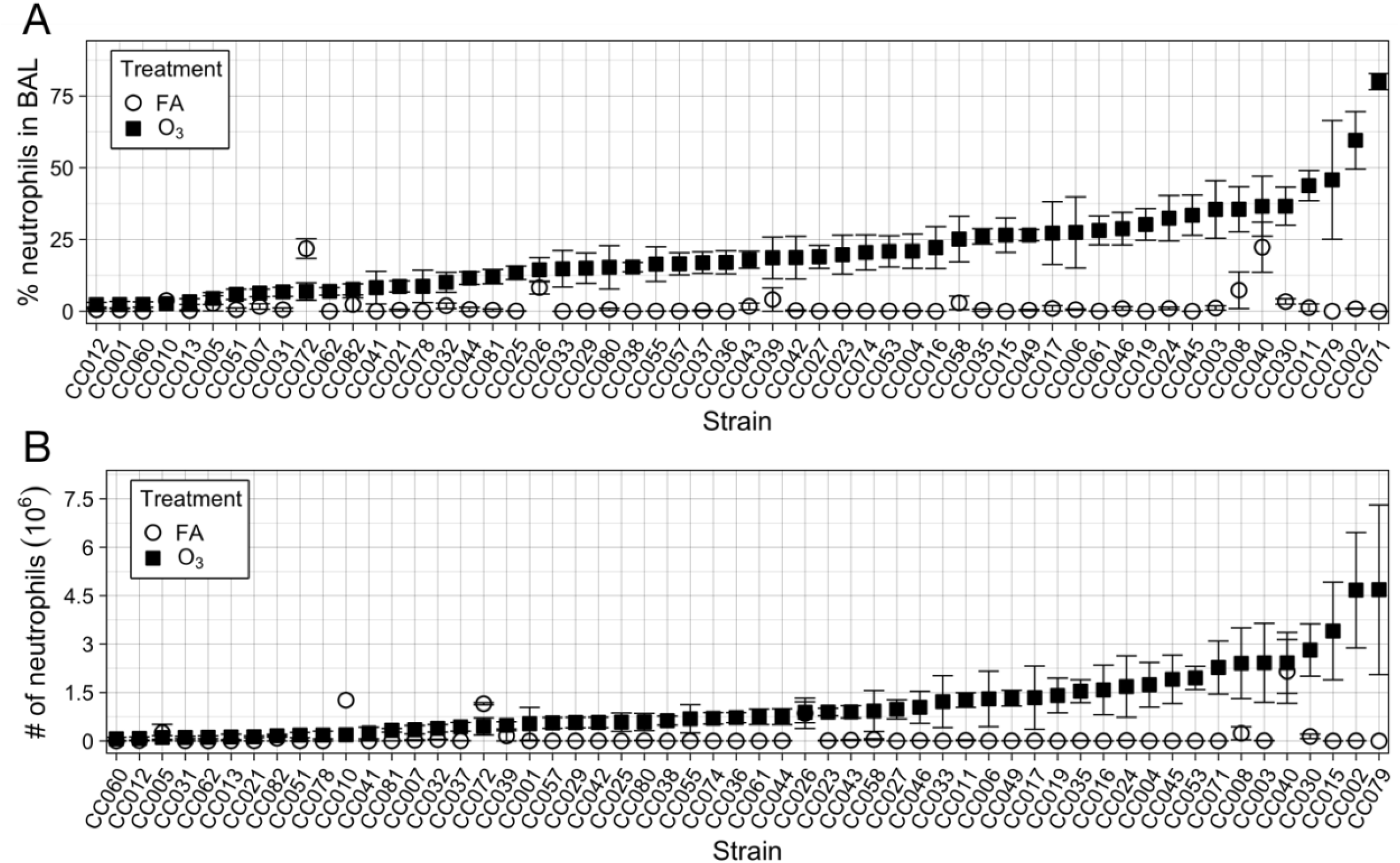
Variation in O_3_-induced neutrophilia across 56 Collaborative Cross (CC) strains. Percent and (B) total number of neutrophils in bronchoalveolar lavage fluid were measured in mice after filtered-air (FA, open circles) or 2 ppm O_3_ exposure (filled squares). Data are represented as strain means and standard error. (n = 4/FA group and 6/O_3_ group, on average)

**Figure 3.**
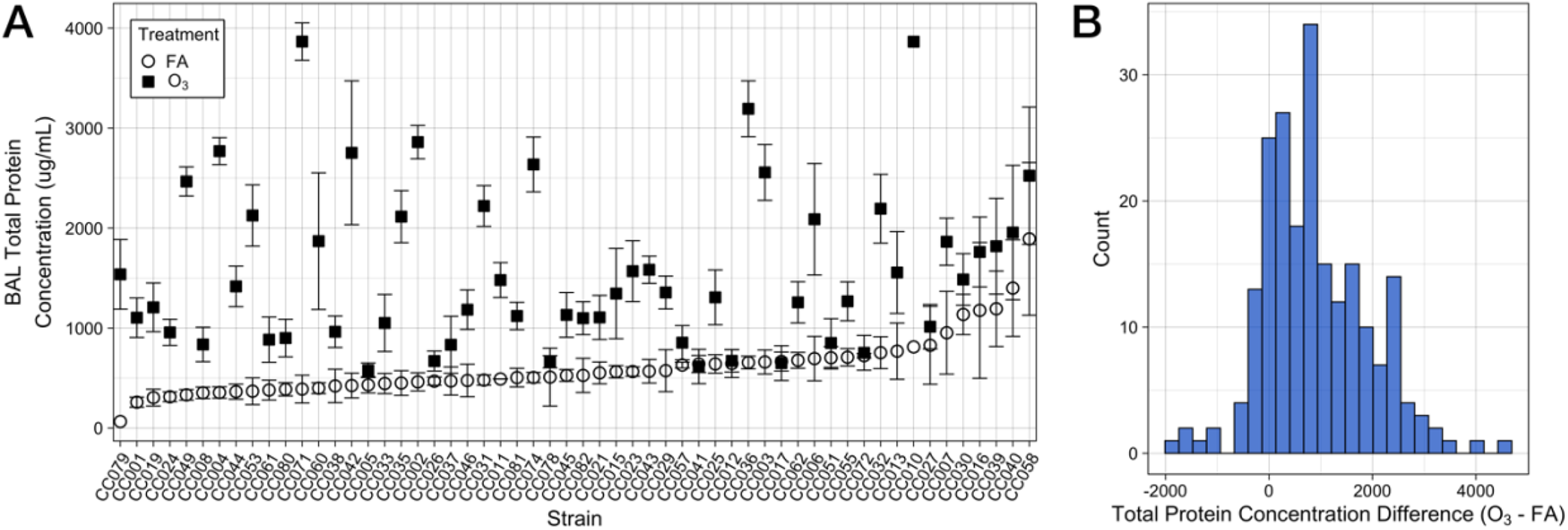
Total protein concentration varies by CC strain and after O_3_ exposure, and protein response is a complex trait. (A) Total protein concentration is modestly influenced by CC strain in control, filtered air (FA)-exposed animals (open circles) and highly variable after O_3_ exposure (closed squares), with indications of strain-by-treatment interactions. (n = 4/FA group and 6/O_3_ group per strain, on average) (B) To examine strain-by-treatment interactions, we defined a protein response phenotype as the difference in total protein concentration between O_3_-exposed mice and FA-exposed mice within a matched pair or trio. The distribution of this phenotype is roughly normally distributed, which suggests that it is a complex trait influenced by multiple genetic loci. (n = 2 pairs and 2 trios per strain, on average)

### Total protein response is associated with genetic variation on Chromosomes 15 and 10

To identify genomic regions associated with variation in O_3_-induced neutrophilia and total protein response, we performed quantitative trait locus (QTL) mapping using a haplotype-based regression procedure accounting for the effects of sex and batch. While we did not detect significant loci associated with neutrophilia, two QTL were associated with total protein response: one locus each on Chromosomes (Chr.) 15 (*Oipq1*, ozone-induced protein response QTL 1) and 10 (*Oipq2)*, the first of which surpassed the genome-wide significance threshold of α = 0.05 (Figure 4A). These loci explained roughly 27 and 13% of variation in the phenotype, respectively.

**Figure 4.**
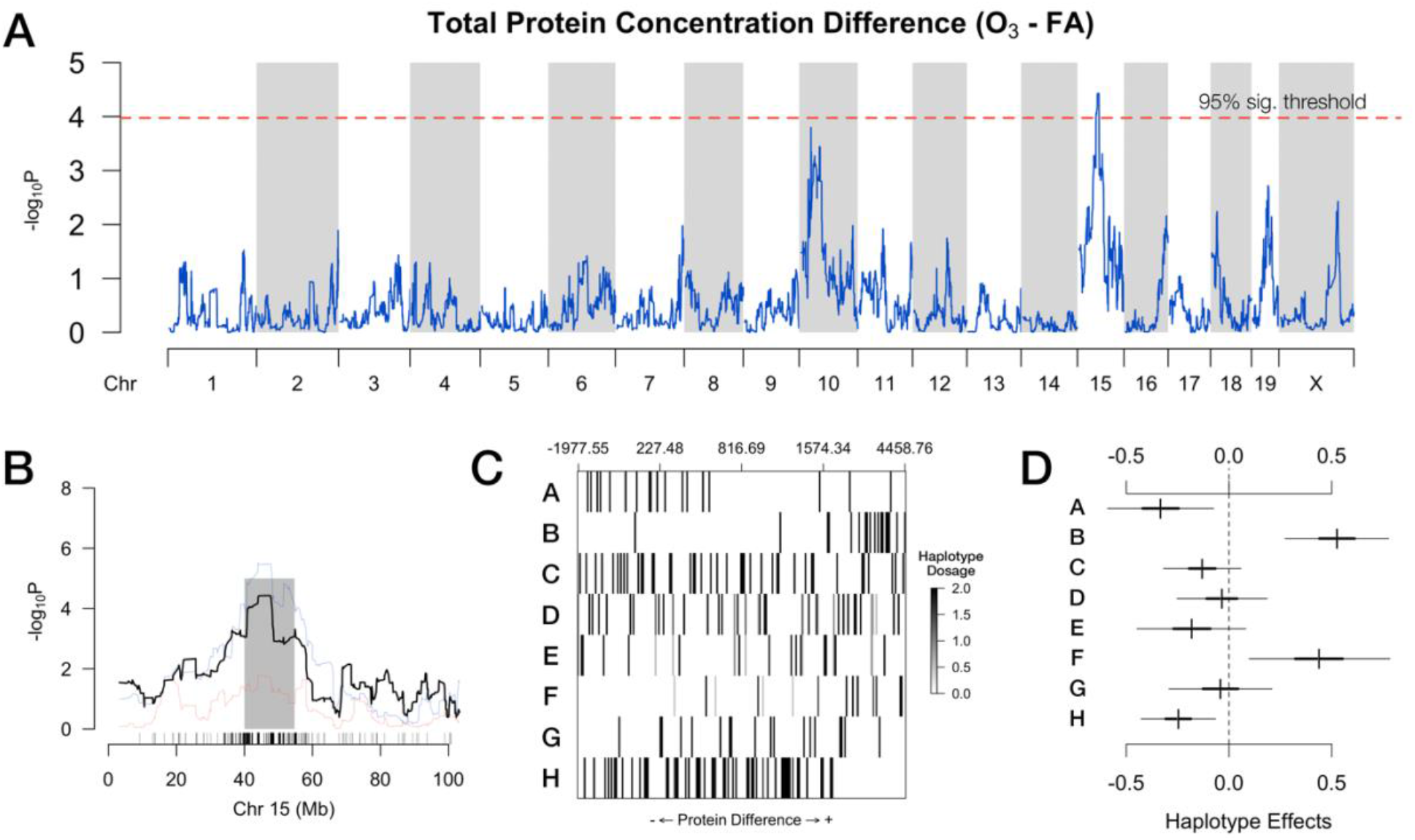
Quantitative trait locus (QTL) mapping uncovers a locus on Chromosome 15 significantly associated with O_3_-induced protein response. (A) QTL scan highlighting regions where variation in total protein concentration difference (i.e., protein response) is associated with haplotype. The red dashed line indicates the 95% significance threshold (red), and one locus on Chromosome (Chr.) 15 surpasses this threshold (marker: JAX00061625_to_UNC25526264, 47.12 Mb). A second locus, on Chromosome 10, surpasses the 90% significance threshold (blue). An 80% confidence interval for the locus on Chr. 15 spans ∼15 Mb (40.17-54.88 Mb), indicated by the gray shading. (C) Haplotype dosage intensities at the Chr. 15 peak marker for strain pairs, ordered by increasing protein response along the x-axis. Darker ticks indicate higher probabilities of a given haplotype at the peak marker. Strain matches with A/J or WSB/EiJ haplotypes are associated with low protein response, while those with C57BL/6J or CAST/EiJ are associated with high protein response. (D) Haplotype effects and confidence intervals at the peak marker within the Chr. 15 QTL. A: A/J, B: C57BL/6J, C: 129S1SvImJ, D: NOD/ShiLtJ, E: NZO/ H1LtJ, F: CAST/EiJ, G: PWK/PhJ, H: WSB/EiJ.

*Oipq1* mapped to a region on Chr. 15 extending ∼15 Mb (80% CI: 40.17-54.88 Mb) which contained 31 protein-coding genes (Figure 4B). To identify candidate genes underlying this QTL, we initially inspected the founder haplotype effects pattern to identify whether there were functionally distinct alleles that could be used to partition and prioritize candidates. Founder haplotype probabilities at the peak marker within *Oipq1* (JAX00061625_to_UNC25526264: 47.12 Mb) showed an overrepresentation of CC strain pairs with C57BL/6J or CAST/EiJ associated with high total protein response, while strain pairs with A/J or WSB/EiJ were associated with low total protein response (Figure 4C). Haplotype substitution effects were modeled using Diploffect (43), indicating a strong positive effect of the C57BL/6J and CAST/EiJ haplotypes on total protein response (Figure 4D). It should be noted that estimated allele effects for CAST/EiJ are less certain than for C57BL/6J due to lower representation of and less confident genotype calls for this founder haplotype at the peak locus (Figure 4C). Interestingly, when visualizing the subspecies origin of the CC founder haplotypes within the QTL interval using the Mouse Phylogeny Viewer (53, 54), we discovered a ∼2 Mb region of *Mus musculus domesticus* intersubspecific introgression into the genome of CAST/EiJ (which is of *castaneus* origin) from ∼43-45 Mb (Supplemental Figure S3). We then used the statistical approach TIMBR (44) to infer the allelic series, *i*.*e*., the number of functional alleles into which the founder haplotypes group. The results indicated a high likelihood of two functional alleles at this QTL, with greatest weight for a haplotype grouping of C57BL/6J and CAST/EiJ (Supplemental Figure S4, Supplemental Tables S1 & S2). We note that, as we observed with Diploffect, there was a similar level of uncertainty for CAST/EiJ haplotype effects (Supplemental Figure S4). Hence, while there is some evidence to suggest that C57BL/6J and CAST/EiJ founder haplotypes share one or more variants within the region that are functionally distinct from other CC founder haplotypes and are causally related to the protein response to O_3_ exposure, this haplotype grouping should be interpreted with caution. Thus, in subsequent analyses, we focused on variants with strain distribution patterns for which C57BL/6J was unique, or C57BL/6J and CAST/EiJ shared a common variant.

*Oipq2* spanned a ∼19 Mb region (80% CI: 24.58-43.61 Mb) encompassing 90 protein-coding genes on Chr. 10 (Supplemental Figure S5A). We examined the founder haplotype probabilities at the peak marker (UNC17621935: 26.204 Mb), which were sorted into a less defined pattern than at *Oipq1*. Strain matches with WSB/EiJ haplotype at this marker were associated with high total protein response, but all other haplotypes were spread evenly across the phenotypic spectrum (Supplemental Figure S5B). Intriguingly, within much of this region (including at the peak marker) in the CC strains surveyed, there is essentially no or low-confidence representation of the PWK/PhJ haplotype, and low representation of the other two wild-derived strains (30). Previous work has established that PWK/PhJ has reduced genome-wide contributions in living CC strains (30), and selection against the PWK/PhJ haplotype may have occurred at this locus to maintain reproductive compatibility in the course of inbreeding (55). Because *Oipq1* surpassed the genome-wide significance threshold and had more clearly defined haplotype effects, we prioritized this locus for further analysis.

### Multiallelic merge analysis identifies multiple candidate variants within Chr. 15 QTL region

To further rank candidate genes and variants within the QTL region, we performed a merge analysis with modifications to accommodate multi-allelic variants, as described previously (45–47). In brief, this approach moves from association at the haplotype level to association at the level of individual variants (both single nucleotides and small insertions/deletions) with variation in the phenotype of interest. By using sequence information from the Inbred Strain Variant Database (56) for each of the CC founders and CC strains that were used for QTL mapping, variants can be identified that are distributed amongst the CC strains in concordance with the haplotype effects pattern. Founder strain haplotypes are “merged” into 2-7 groups in accordance with how the variants are distributed (*i*.*e*., the strain distribution profile). Variants in the merged model that explain trait variation equally well or better than the full haplotype model (but with fewer parameters) can be considered candidate quantitative trait variants (QTVs). In the *Oipq1* region, 995 variants were identified using this method, with a -log_10_(*p*-value) > 4 (roughly the cut-off used for QTL mapping; Figure 5A). Variants identified using merge analysis represented a variety of strain distribution patterns (SDPs), with two SDPs more common than the others: C57BL/6J alone or C57BL/6J and CAST/EiJ discordant from all other strains (Figure 5B). These variants were predicted to have several consequences on gene/protein function, though most were present within introns or other non-coding regions of the QTL (Figure 5C). Overall, this subset of variants was located within or near 21 genes (13 protein-coding, 8 predicted), most of which were concentrated within the region between ∼40.9-44.6 Mb containing 16 genes (10 protein-coding) (Figure 5D, Supplemental Table S4).

**Figure 5.**
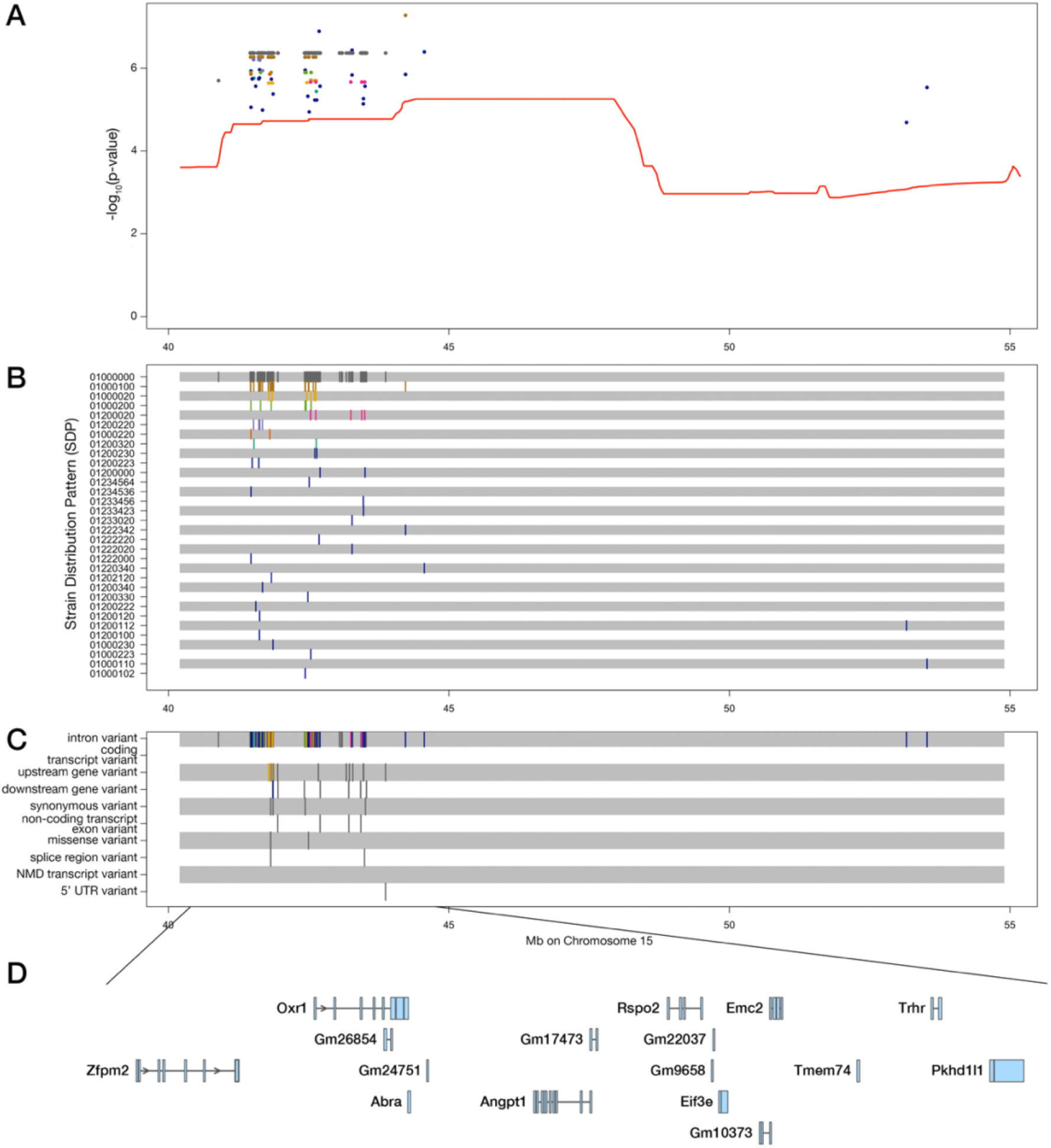
Candidate variant (merge) analysis pinpoints variants and narrows the list of genes of interest in the Chr. 15 QTL. (A) Merge analysis identified 995 variants whose strain distribution pattern (SDP) explains the variation in protein response with a more parsimonious fit than the 8-haplotype model used for QTL mapping (indicated by the red line). (B) Listed in the left margin of the lower tracks are the recoded allelic series for each SDP. For example, the allelic series of a biallelic variant is comprised by 0s and 1s. Thus, the first track is a biallelic variant where all seven founder strains share the same allele (0) and C57BL/6J has the alternative allele (1). (C) Corresponding consequences for each of the variants present in (A) and(B), where color-coding is consistent across both panels. Note that variants may have more than one consequence, and thus may appear across multiple tracks. (D) The 16 genes present within the genomic interval with the highest density of variants (∼41-45 Mb). Of these genes, 10 are protein-coding.

We used a weight of evidence approach to further prioritize candidate genes within the interval, using the following criteria: (1) presence of variants that are concordant with the haplotype effects pattern, with an emphasis on C57BL/6J and/or CAST/EiJ discordant from all other founder strains; (2) presence of coding variants that lead to amino acid alterations; (3) evidence that a gene is expressed the lungs (from our previously unpublished and published data (39), or publicly available datasets); and (4) biological relevance, as judged by a gene’s known function and prior description in the literature (Table 1). Two genes within the interval met all criteria: *Angpt1* (angiopoietin-1) and *Oxr1* (oxidation resistance 1). A third candidate (*Rspo2*, R-spondin 2) met three of the criteria used, and extensive prior annotation in the literature provided additional biological plausibility.

**Table 1.**
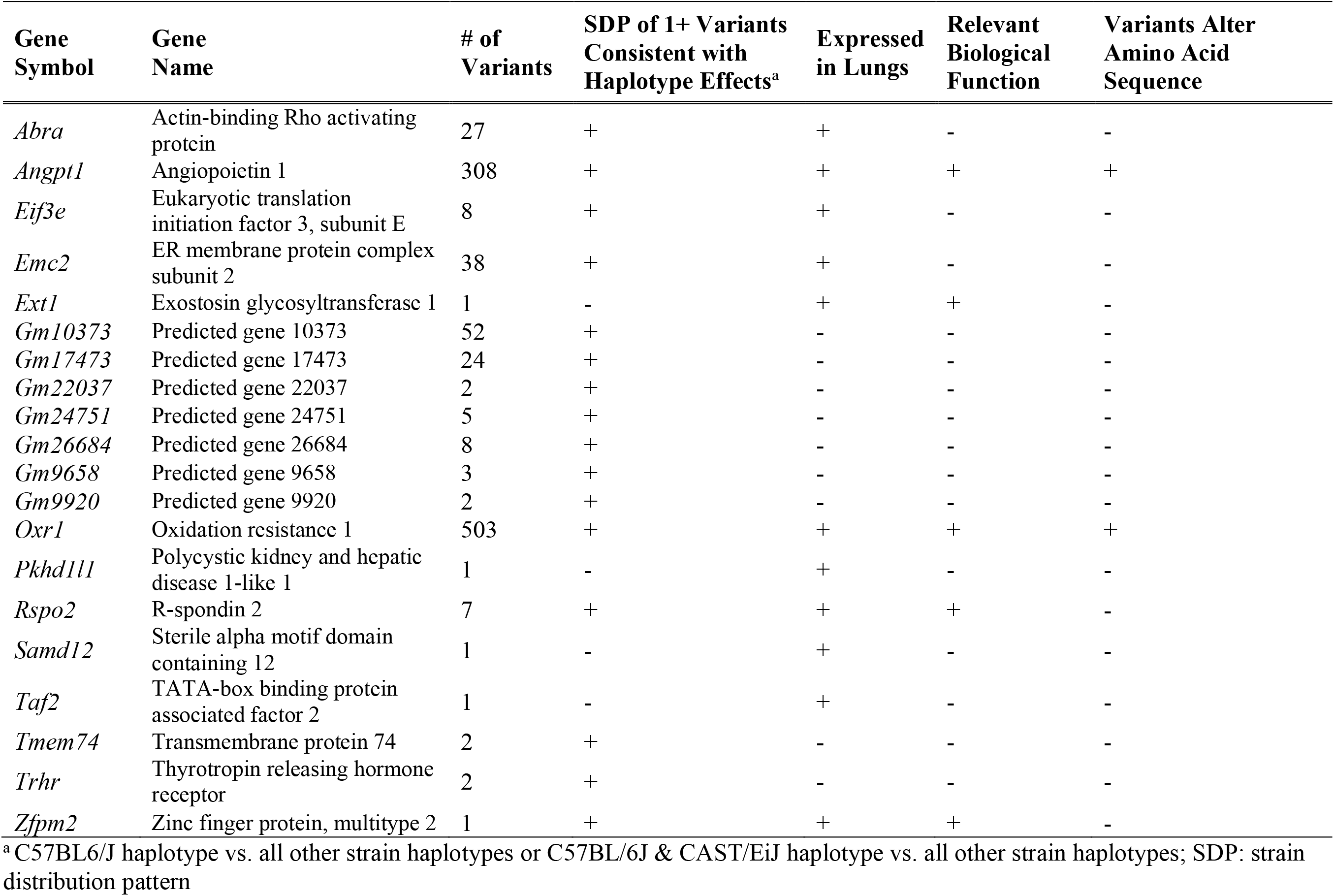
Weight of evidence analysis in *Oipq1* candidate genes.

Using information from Ensembl, UniProt, and multiple variant consequence prediction tools, we inspected the three missense variants within *Oxr1* (rs50179186, rs31574788) and *Angpt1* (rs32511504) to determine whether any had putative consequences on protein structure and/or function (Table 2 & Supplemental Table S4). All three variants had SDPs where C57BL/6J was distinct from the other founder strains. Both variants within *Oxr1* were present within a predicted disordered region, while the variant within *Angpt1* was in a linker region, between coiled-coiled and fibrinogen domains. This analysis was largely inconclusive for two of the three variants (rs50179186 in *Oxr1* and rs32511504 in *Angpt1*), as the results across tools were conflicting which has been observed in many previous investigations (51, 57). The second variant in *Oxr1* (rs31574788) appeared to have no predicted effect on protein function across all four tools.

**Table 2.**
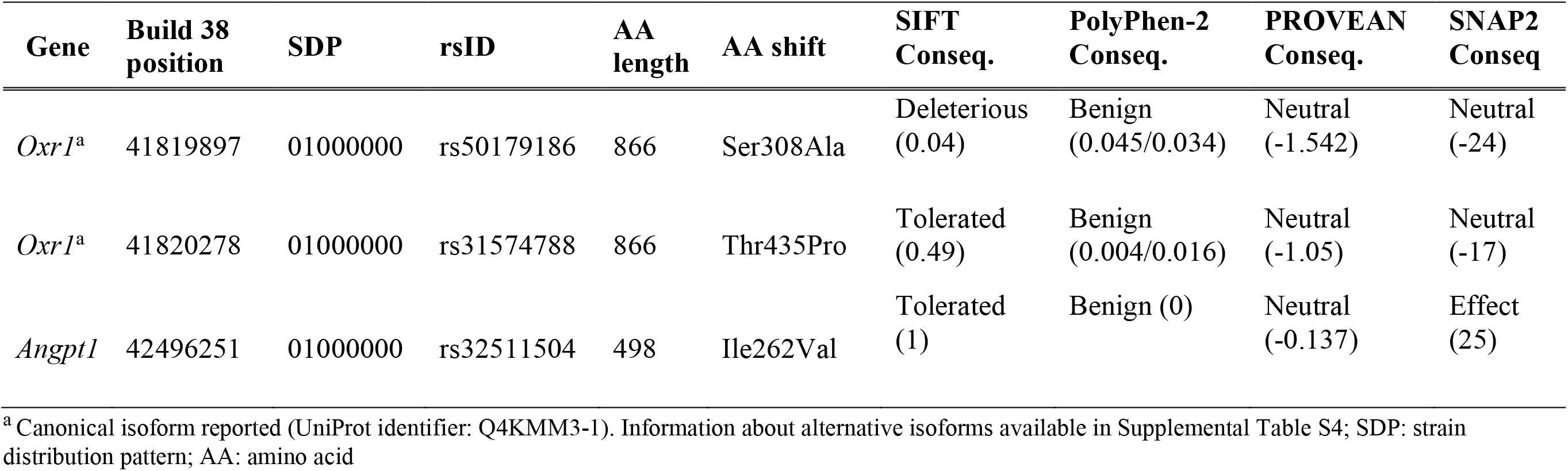
Table of missense variants in candidate genes within the *Oipq1* region. Scores for SIFT, PolyPhen-2, PROVEAN, and SNAP2 are present in parentheses next to their predicted consequences.

## Discussion

Here, we present discovery of two loci associated with ozone (O_3_)-induced lung injury, *Oipq1* and *Oipq2*, located on Chr. 15 and 10, respectively. Notably, the region encompassed by *Oipq1* has been previously implicated in two studies aiming to identify genetic variants associated with lung injury. One study utilized a C57BL/6J:129X1/SvJ F2 population to map QTL driving fatality due to hyperoxic (58). In that study, the C57BL/6J allele at the QTL was associated with higher lung injury, consistent with the results of our study. Similarly, an overlapping QTL for pulmonary hemorrhage was identified in a study utilizing a CC003/Unc:CC053/Unc F2 intercross to identify genetic determinants of pulmonary responses to SARS-CoV. Though this QTL was not the focus of their study (59), the authors did observe a positive association between the CAST/EiJ allele and pulmonary hemorrhage and the inverse relationship with the PWK/PhJ allele, matching the haplotype effects observed in our study. Therefore, there is strong evidence that this locus mediates responses to multiple stimuli that induce lung injury.

Within *Oipq1*, we nominated three candidate genes: *Rspo2* (R-spondin 2), *Angpt1* (angiopoietin-1), and *Oxr1* (oxidation resistance 1). Information about missense variants within *Angpt1* and *Oxr1* is lacking, because the results from variant prediction tools were conflicting and/or largely suggested that the missense variants had neutral effects on protein function. While we used a suite of approaches that each incorporate unique features for prediction including sequence homology, physico-chemical properties, and predicted secondary structure, even state-of-the-art methods fail to fully recapitulate functional characterization through experimental strategies (*i*.*e*., deep mutational scanning), which remain the gold standard (60). Thus, direct experimental classification is needed to determine whether these variants have effects on protein function. It is also possible, perhaps even likely, that variants in this region affect the expression of these candidate genes (*i*.*e*., are eQTL), and that variation in gene expression underlies the *<u>Oipq1</u>* QTL. Future studies addressing this hypothesis are also needed.

One candidate gene at *Oipq1, Rspo2*, is a member of the R-spondin (RSPO) protein family, a class of secreted ligands known to regulate Wnt signaling, tissue regeneration and organization in various regions of the body (61). In particular, RSPO2 is required for lung, limb, and craniofacial development (62), and *Rspo2*-deficient mice are born with various skeletal defects (63, 64), and die immediately upon birth due to respiratory failure (65, 66). Recessive mutations in *RSPO2* cause tetra-amelia syndrome-2, a human syndrome characterized by partial or complete absence of limbs along with incomplete lung development (67). Jackson, *et al*. recently examined the consequences of *Rspo2* conditional deletion in adult mice. They reported that *Rspo2* loss caused lung neutrophilia via a disrupted lung endothelial barrier (68). While there are no variants within this gene, we noted multiple noncoding variants within the in/near the gene, and its function is in alignment with features of the pathology caused by O_3_ exposure. Its expression is largely restricted to the lung mesenchyme; therefore, we were unable to directly interrogate whether it is altered by O_3_ exposure using our previously published data or other datasets. Additionally, its expression is quite low in all life stages beyond embryonic development, thus detecting whether *Rspo2* expression is changed in the context of other lung insults using publicly available data has proven challenging. Nevertheless, the phenotypes observed after its conditional deletion in adulthood are compelling, and warrant further investigation in the context of O_3_-induced lung injury.

The second candidate gene *Angpt1* encodes angiopoietin-1 (ANGPT1), which is a secreted ligand that binds to and activates the Tie2 receptor tyrosine kinase (encoded by *Tek*) to promote endothelial barrier function and vascular growth (69). It has also been suggested that ANGPT1 activity can improve and reinforce leaky or otherwise poorly functioning vessels (70, 71). ANGPT1 is largely produced by vascular support cells and platelets, and its activity can be antagonized by ANGPT2 (72). Disrupted balance of ANGPT2/ANGPT1 in serum is associated with poor outcomes in a variety of disease states, including acute respiratory distress syndrome, bronchopulmonary dysplasia, and sepsis (73–75). Genetic association studies have identified variants in *ANGPT2* associated with susceptibility to acute lung injury (76). Together, these studies suggest that ANGPT2 is often a pathogenic regulator of acute lung injury and barrier dysfunction, and balance to this system can be restored by supplementation with ANGPT1. Thus, one hypothesis arising from our study is that in strains with haplotypes associated with more severe protein response (*i*.*e*., C57BL/6J and potentially CAST/EiJ), there may be variants in or near *Angpt1* that alter its activity and/or function, thereby disrupting its ability to negatively regulate ANGPT2/TEK signaling. *In vitro* and *in vivo* functional validation studies in models of O_3_ exposure or other types of acute lung injury will be necessary to evaluate this hypothesis.

Our final candidate gene, *Oxr1* (oxidation resistance 1), has less evidence tying its functions to acute lung injury and epithelial barrier integrity, though these remain to be examined. This protein, while aptly named for a role in response to oxidant gases, has largely been implicated in maintaining genome integrity and cell survival in the face of stressors that cause either oxidative stress-dependent or -independent DNA damage (*e*.*g*., reactive oxygen species, radiation, alkylating agents) (77). This gene was first discovered in an *E. coli* screen for human genes involved in repairing or preventing oxidative DNA damage (78), and later discovered to function largely in the mitochondria (79). *Oxr1* has since been studied largely in the context of neurological diseases including amyotrophic lateral sclerosis (ALS) (80–82) and other neurodegenerative conditions, as well as a recent study describing recessive loss-of-function variants in *OXR1* associated with cerebellar atrophy, seizures, developmental delays, among other clinical features (83). Only a few studies have mentioned this gene in the context of lung disease, including one examining vanadium pentoxide (V_2_O_5_)-induced occupational bronchitis where *OXR1* expression was induced in human lung fibroblasts after exposure to V_2_O_5_(84). It is worth mentioning that mice express multiple isoforms of *Oxr1* whose tissue specificity was recently characterized (85): many tissues expressed the shortest version (*Oxr1D*) and *Oxr1B1*-*4*, while the longest (*Oxr1A*) was restricted to the brain. The cited study went on to characterize the functions of the OXR1A isoform, demonstrating that its TLDc domain (present in all OXR1 isoforms) facilitates interactions with a variety of proteins including the PRMT5 methyltransferase, thus representing one pathway by which this gene alters cellular function in response to stress. However, further studies will be required to understand its roles in oxidative stress responses in the lungs and assess its relationship to the protein response phenotype measured here.

We did not detect any loci associated with variation in airway neutrophilia, another hallmark O_3_ -induced phenotype, despite the fact that others have previously identified QTL for this trait (20). However, we note that this phenotype was highly heritable (*H*^*2*^: ∼0.47 for percentage and ∼0.52 for total number). Thus, one potential explanation for the lack of QTL is that the genetic architecture of airway neutrophilia may be more complex than lung injury and involve contributions from many loci with individually small effects. One option to address this would be to incorporate information about phenotypes that lie along the causal chain from O_3_ exposure to airway neutrophilia (*e*.*g*., cytokines), as we expect these intermediate phenotypes to have higher heritabilities and simpler genetic architectures. Alternatively, to achieve greater statistical power for QTL mapping, one could make use of the Diversity Outbred mouse population since a larger number of genetically distinct mice can be examined in that population (86). However, we note here that the protein response QTL we identified here was detected using a “delta” framework (in which the phenotype was the difference between O_3_- vs FA-exposed mice), which required inbred strains so that baseline (filtered air) effects could be accounted for. In conclusion, we have identified a significant QTL on mouse Chr. 15 associated with O_3_ -induced lung injury. Through additional genetic and bioinformatic analyses, we delimited the QTL region and identified three high priority candidate genes worthy of additional investigation. Our study also demonstrates the utility of the Collaborative Cross genetic reference population for identifying interactions between genetic variants and environmental exposures.

## Acknowledgements

The authors would like to acknowledge the assistance of Daniel Vargas and Jessica Bustamante (logistical support); Courtney Nesline and the UNC Division of Comparative Medicine; Darla Miller, Ginger Shaw, and Dr. Rachel Lynch of the UNC Systems Genetics Core Facility (Collaborative Cross mice); Drs. Greg Keele and Wes Crouse (maintenance of and guidance with using the miqtl and TIMBR R packages, respectively); and Drs. Will Valdar and Yanwei Cai (QTL mapping). This research was funded by NIH Grants ES024965 and ES024965-S1 to S.N.P.K., a T32 training grant (ES007126-35) and a Leon and Bertha Golberg Postdoctoral Fellowship from the UNC Curriculum in Toxicology and Environmental Medicine to G.J.S., and a UNC Dissertation Completion Fellowship to A.T.

## REFERENCES

1. C. S. Kim, et al., Lung function and inflammatory responses in healthy young adults exposed to 0.06 ppm ozone for 6.6 hours. Am. J. Respir. Crit. Care Med. 183, 1215–1221 (2011).

2. E. S. Schelegle, C. A. Morales, W. F. Walby, S. Marion, R. P. Allen, 6.6-hour inhalation of ozone concentrations from 60 to 87 parts per billion in healthy humans. Am. J. Respir. Crit. Care Med. 180, 265–272 (2009).

3. R. M. Aris, et al., Ozone-induced airway inflammation in human subjects as determined by airway lavage and biopsy. Am. Rev. Respir. Dis. 148, 1363–1372 (1993).

4. H. S. Koren, et al., Ozone-induced inflammation in the lower airways of human subjects. Am. Rev. Respir. Dis. 139, 407–415 (1989).

5. R. B. Devlin, et al., Inflammation and cell damage induced by repeated exposure of humans to ozone. Inhal Toxicol 9, 211–235 (1997).

6. H. R. Kehrl, et al., Ozone exposure increases respiratory epithelial permeability in humans. Am. Rev. Respir. Dis. 135, 1124–1128 (1987).

7. R. T. Burnett, J. R. Brook, W. T. Yung, R. E. Dales, D. Krewski, Association between ozone and hospitalization for respiratory diseases in 16 Canadian cities. Environ. Res. 72, 24–31 (1997).

8. J. F. Gent, et al., Association of low-level ozone and fine particles with respiratory symptoms in children with asthma. JAMA 290, 1859–1867 (2003).

9. M. J. Strickland, et al., Short-term associations between ambient air pollutants and pediatric asthma emergency department visits. Am. J. Respir. Crit. Care Med. 182, 307–316 (2010).

10. L. A. Darrow, et al., Air pollution and acute respiratory infections among children 0-4 years of age: an 18-year time-series study. Am. J. Epidemiol. 180, 968–977 (2014).

11. J. Ciencewicki, S. Trivedi, S. R. Kleeberger, Oxidants and the pathogenesis of lung diseases. J. Allergy Clin. Immunol. 122, 456–68; quiz 469–70 (2008).

12. G. D. Thurston, et al., Outdoor Air Pollution and New-Onset Airway Disease. An Official American Thoracic Society Workshop Report. Ann. Am. Thorac. Soc. 17, 387–398 (2020).

13. L. G. Que, J. V. Stiles, J. S. Sundy, W. M. Foster, Pulmonary function, bronchial reactivity, and epithelial permeability are response phenotypes to ozone and develop differentially in healthy humans. J. Appl. Physiol. 111, 679–687 (2011).

14. O. Holz, et al., Ozone-induced airway inflammatory changes differ between individuals and are reproducible. Am. J. Respir. Crit. Care Med. 159, 776–784 (1999).

15. W. F. McDonnell 3rd, D. H. Horstman, S. Abdul-Salaam, D. E. House, Reproducibility of individual responses to ozone exposure. Am. Rev. Respir. Dis. 131, 36–40 (1985).

16. N. E. Alexis, et al., The glutathione-S-transferase Mu 1 null genotype modulates ozone-induced airway inflammation in human subjects. J. Allergy Clin. Immunol. 124, 1222-1228.e5 (2009).

17. I. A. Yang, et al., Association of tumor necrosis factor-alpha polymorphisms and ozone-induced change in lung function. Am. J. Respir. Crit. Care Med. 171, 171–176 (2005).

18. S. E. Alexeeff, et al., Ozone exposure, antioxidant genes, and lung function in an elderly cohort: VA normative aging study. Occup. Environ. Med. 65, 736–742 (2008).

19. H. Moreno-Macías, et al., Ozone exposure, vitamin C intake, and genetic susceptibility of asthmatic children in Mexico City: a cohort study. Respir. Res. 14, 14 (2013).

20. S. R. Kleeberger, et al., Linkage analysis of susceptibility to ozone-induced lung inflammation in inbred mice. Nat. Genet. 17, 475–478 (1997).

21. D. R. Prows, H. G. Shertzer, M. J. Daly, C. L. Sidman, G. D. Leikauf, Genetic analysis of ozone-induced acute lung injury in sensitive and resistant strains of mice. Nat. Genet. 17, 471–474 (1997).

22. S. R. Kleeberger, S. Reddy, L. Y. Zhang, A. E. Jedlicka, Genetic susceptibility to ozone-induced lung hyperpermeability: role of toll-like receptor 4. Am. J. Respir. Cell Mol. Biol. 22, 620–627 (2000).

23. H. Y. Cho, L. Y. Zhang, S. R. Kleeberger, Ozone-induced lung inflammation and hyperreactivity are mediated via tumor necrosis factor-alpha receptors. Am. J. Physiol. Lung Cell. Mol. Physiol. 280, L537–46 (2001).

24. A. K. Bauer, et al., Identification of candidate genes downstream of TLR4 signaling after ozone exposure in mice: a role for heat-shock protein 70. Environ. Health Perspect. 119, 1091–1097 (2011).

25. L. Fakhrzadeh, J. D. Laskin, D. L. Laskin, Deficiency in inducible nitric oxide synthase protects mice from ozone-induced lung inflammation and tissue injury. Am. J. Respir. Cell Mol. Biol. 26, 413–419 (2002).

26. A. J. Connor, J. D. Laskin, D. L. Laskin, Ozone-induced lung injury and sterile inflammation. Role of toll-like receptor 4. Exp. Mol. Pathol. 92, 229–235 (2012).

27. S. A. Shore, I. N. Schwartzman, B. Le Blanc, G. G. Murthy, C. M. Doerschuk, Tumor necrosis factor receptor 2 contributes to ozone-induced airway hyperresponsiveness in mice. Am. J. Respir. Crit. Care Med. 164, 602–607 (2001).

28. G. A. Churchill, et al., The Collaborative Cross, a community resource for the genetic analysis of complex traits. Nat. Genet. 36, 1133–1137 (2004).

29. Collaborative Cross Consortium, The genome architecture of the Collaborative Cross mouse genetic reference population. Genetics 190, 389–401 (2012).

30. A. Srivastava, et al., Genomes of the Mouse Collaborative Cross. Genetics 206, 537–556 (2017).

31. D. L. Aylor, et al., Genetic analysis of complex traits in the emerging Collaborative Cross. Genome Res. 21, 1213–1222 (2011).

32. V. M. Philip, et al., Genetic analysis in the Collaborative Cross breeding population. Genome Res. 21, 1223–1238 (2011).

33. S. N. Kelada, et al., Integrative genetic analysis of allergic inflammation in the murine lung. Am. J. Respir. Cell Mol. Biol. 51, 436–445 (2014).

34. L. J. Donoghue, et al., Identification of trans Protein QTL for Secreted Airway Mucins in Mice and a Causal Role for Bpifb1. Genetics 207, 801–812 (2017).

35. H. Rutledge, et al., Genetic regulation of Zfp30, CXCL1, and neutrophilic inflammation in murine lung. Genetics 198, 735–745 (2014).

36. M. T. Ferris, et al., Modeling host genetic regulation of influenza pathogenesis in the collaborative cross. PLoS Pathog. 9, e1003196 (2013).

37. L. E. Gralinski, et al., Genome Wide Identification of SARS-CoV Susceptibility Loci Using the Collaborative Cross. PLoS Genet. 11, e1005504 (2015).

38. G. J. Smith, L. Walsh, M. Higuchi, S. N. P. Kelada, Development of a large-scale computer-controlled ozone inhalation exposure system for rodents. Inhal. Toxicol. 31, 61– 72 (2019).

39. A. Tovar, et al., Transcriptional Profiling of the Murine Airway Response to Acute Ozone Exposure. Toxicol. Sci. 173, 114–130 (2020).

40. A. M. Holand, I. Steinsland, S. Martino, H. Jensen, Animal models and integrated nested Laplace approximations. G3 3, 1241–1251 (2013).

41. G. R. Keele, et al., Genetic Fine-Mapping and Identification of Candidate Genes and Variants for Adiposity Traits in Outbred Rats. Obesity 26, 213–222 (2018).

42. G. R. Keele, et al., Integrative QTL analysis of gene expression and chromatin accessibility identifies multi-tissue patterns of genetic regulation. PLoS Genet. 16, e1008537 (2020).

43. Z. Zhang, W. Wang, W. Valdar, Bayesian modeling of haplotype effects in multiparent populations. Genetics 198, 139–156 (2014).

44. W. L. Crouse, S. N. P. Kelada, W. Valdar, Inferring the Allelic Series at QTL in Multiparental Populations. Genetics 216, 957–983 (2020).

45. M. Mosedale, et al., Identification of Candidate Risk Factor Genes for Human Idelalisib Toxicity Using a Collaborative Cross Approach. Toxicol. Sci. 172, 265–278 (2019).

46. M. Mosedale, et al., Editor’s Highlight: Candidate Risk Factors and Mechanisms for Tolvaptan-Induced Liver Injury Are Identified Using a Collaborative Cross Approach. Toxicol. Sci. 156, 438–454 (2017).

47. B. Yalcin, J. Flint, R. Mott, Using progenitor strain information to identify quantitative trait nucleotides in outbred mice. Genetics 171, 673–681 (2005).

48. P. C. Ng, S. Henikoff, Predicting deleterious amino acid substitutions. Genome Res. 11, 863–874 (2001).

49. I. A. Adzhubei, et al., A method and server for predicting damaging missense mutations. Nat. Methods 7, 248–249 (2010).

50. Y. Choi, G. E. Sims, S. Murphy, J. R. Miller, A. P. Chan, Predicting the functional effect of amino acid substitutions and indels. PLoS One 7, e46688 (2012).

51. M. Hecht, Y. Bromberg, B. Rost, Better prediction of functional effects for sequence variants. BMC Genomics 16 Suppl 8, S1 (2015).

52. K. Gabehart, K. A. Correll, J. E. Loader, C. W. White, A. Dakhama, The lung response to ozone is determined by age and is partially dependent on toll-Like receptor 4. Respir. Res. 16, 117 (2015).

53. H. Yang, et al., Subspecific origin and haplotype diversity in the laboratory mouse. Nat. Genet. 43, 648–655 (2011).

54. J. R. Wang, F. P.-M. de Villena, L. McMillan, Comparative analysis and visualization of multiple collinear genomes. BMC Bioinformatics 13 Suppl 3, S13 (2012).

55. J. R. Shorter, et al., Male Infertility Is Responsible for Nearly Half of the Extinction Observed in the Mouse Collaborative Cross. Genetics 206, 557–572 (2017).

56. D. Oreper, Y. Cai, L. M. Tarantino, F. P.-M. de Villena, W. Valdar, Inbred Strain Variant Database (ISVdb): A Repository for Probabilistically Informed Sequence Differences Among the Collaborative Cross Strains and Their Founders. G3 7, 1623–1630 (2017).

57. C. Dong, et al., Comparison and integration of deleteriousness prediction methods for nonsynonymous SNVs in whole exome sequencing studies. Hum. Mol. Genet. 24, 2125– 2137 (2015).

58. D. R. Prows, et al., Genetic analysis of hyperoxic acute lung injury survival in reciprocal intercross mice. Physiol. Genomics 30, 271–281 (2007).

59. L. E. Gralinski, et al., Allelic Variation in the Toll-Like Receptor Adaptor Protein Ticam2 Contributes to SARS-Coronavirus Pathogenesis in Mice. G3 7, 1653–1663 (2017).

60. J. Reeb, T. Wirth, B. Rost, Variant effect predictions capture some aspects of deep mutational scanning experiments. BMC Bioinformatics 21, 107 (2020).

61. W. B. M. de Lau, B. Snel, H. C. Clevers, The R-spondin protein family. Genome Biol. 13, 242 (2012).

62. S. M. Bell, et al., R-spondin 2 is required for normal laryngeal-tracheal, lung and limb morphogenesis. Development 135, 1049–1058 (2008).

63. S. M. Bell, C. M. Schreiner, K. A. Hess, K. P. Anderson, W. J. Scott, Asymmetric limb malformations in a new transgene insertional mutant, footless. Mech. Dev. 120, 597–605 (2003).

64. M. Aoki, H. Kiyonari, H. Nakamura, H. Okamoto, R-spondin2 expression in the apical ectodermal ridge is essential for outgrowth and patterning in mouse limb development. Dev. Growth Differ. 50, 85–95 (2008).

65. W. Yamada, et al., Craniofacial malformation in R-spondin2 knockout mice. Biochem. Biophys. Res. Commun. 381, 453–458 (2009).

66. J.-S. Nam, et al., Mouse R-spondin2 is required for apical ectodermal ridge maintenance in the hindlimb. Dev. Biol. 311, 124–135 (2007).

67. E. Szenker-Ravi, et al., RSPO2 inhibition of RNF43 and ZNRF3 governs limb development independently of LGR4/5/6. Nature 557, 564–569 (2018).

68. S. R. Jackson, et al., R-spondin 2 mediates neutrophil egress into the alveolar space through increased lung permeability. BMC Res. Notes 13, 54 (2020).

69. C. Suri, et al., Requisite role of angiopoietin-1, a ligand for the TIE2 receptor, during embryonic angiogenesis. Cell 87, 1171–1180 (1996).

70. G. Thurston, et al., Leakage-resistant blood vessels in mice transgenically overexpressing angiopoietin-1. Science 286, 2511–2514 (1999).

71. G. Thurston, et al., Angiopoietin-1 protects the adult vasculature against plasma leakage. Nat. Med. 6, 460–463 (2000).

72. P. C. Maisonpierre, et al., Angiopoietin-2, a natural antagonist for Tie2 that disrupts in vivo angiogenesis. Science 277, 55–60 (1997).

73. S. M. Parikh, et al., Excess circulating angiopoietin-2 may contribute to pulmonary vascular leak in sepsis in humans. PLoS Med. 3, e46 (2006).

74. V. Bhandari, et al., Hyperoxia causes angiopoietin 2-mediated acute lung injury and necrotic cell death. Nat. Med. 12, 1286–1293 (2006).

75. D. C. Gallagher, et al., Circulating angiopoietin 2 correlates with mortality in a surgical population with acute lung injury/adult respiratory distress syndrome. Shock 29, 656–661 (2008).

76. N. J. Meyer, et al., ANGPT2 genetic variant is associated with trauma-associated acute lung injury and altered plasma angiopoietin-2 isoform ratio. Am. J. Respir. Crit. Care Med. 183, 1344–1353 (2011).

77. A. Matsui, K. Hashiguchi, M. Suzuki, Q.-M. Zhang-Akiyama, Oxidation resistance 1 functions in the maintenance of cellular survival and genome stability in response to oxidative stress-independent DNA damage. Genes Environ 42, 29 (2020).

78. M. R. Volkert, N. A. Elliott, D. E. Housman, Functional genomics reveals a family of eukaryotic oxidation protection genes. Proc. Natl. Acad. Sci. U. S. A. 97, 14530–14535 (2000).

79. N. A. Elliott, M. R. Volkert, Stress induction and mitochondrial localization of Oxr1 proteins in yeast and humans. Mol. Cell. Biol. 24, 3180–3187 (2004).

80. P. L. Oliver, et al., Oxr1 is essential for protection against oxidative stress-induced neurodegeneration. PLoS Genet. 7, e1002338 (2011).

81. K. X. Liu, et al., Neuron-specific antioxidant OXR1 extends survival of a mouse model of amyotrophic lateral sclerosis. Brain 138, 1167–1181 (2015).

82. M. J. Finelli, K. X. Liu, Y. Wu, P. L. Oliver, K. E. Davies, Oxr1 improves pathogenic cellular features of ALS-associated FUS and TDP-43 mutations. Hum. Mol. Genet. 24, 3529–3544 (2015).

83. J. Wang, et al., Loss of Oxidation Resistance 1, OXR1, Is Associated with an Autosomal-Recessive Neurological Disease with Cerebellar Atrophy and Lysosomal Dysfunction. Am. J. Hum. Genet. 105, 1237–1253 (2019).

84. J. L. Ingram, et al., Genomic analysis of human lung fibroblasts exposed to vanadium pentoxide to identify candidate genes for occupational bronchitis. Respir. Res. 8, 34 (2007).

85. M. Yang, et al., OXR1A, a Coactivator of PRMT5 Regulating Histone Arginine Methylation. Cell Rep. 30, 4165-4178.e7 (2020).

86. K. L. Svenson, et al., High-resolution genetic mapping using the Mouse Diversity outbred population. Genetics 190, 437–447 (2012).

