## Supplemental Figures for "A novel gene-by-environment quantitative trait locus on mouse chromosome 15 underlies susceptibility to acute ozone-induced lung injury" (from another version of this preprint: version 1, 2021-05-20)

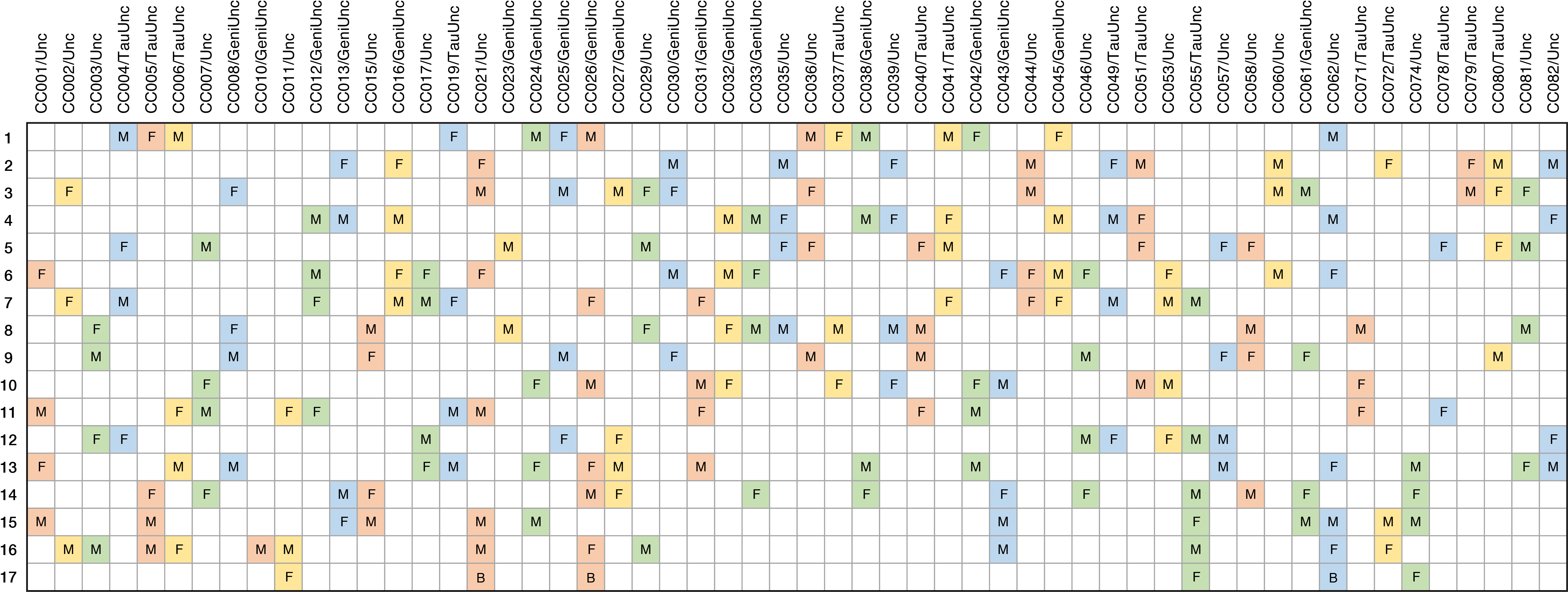

**Supplemental Figure S1. Calendar of experimental batches.** M: male, F: female, B: both sexes represented; Batch 1: 10/2/18, Batch 2: 10/4/18, Batch 3: 10/9/18, Batch 4: 10/11/18, Batch 5: 10/17/18, Batch 6: 10/19/18, Batch 7: 10/23/18, Batch 8: 10/25/18, Batch 9: 10/29/18, Batch 10: 11/1/18, Batch 11: 11/6/18, Batch 12: 11/8/18, Batch 13: 11/16/18, Batch 14: 11/20/18, Batch 15: 11/27/18, Batch 16: 12/5/18, Batch 17: 12/11/1

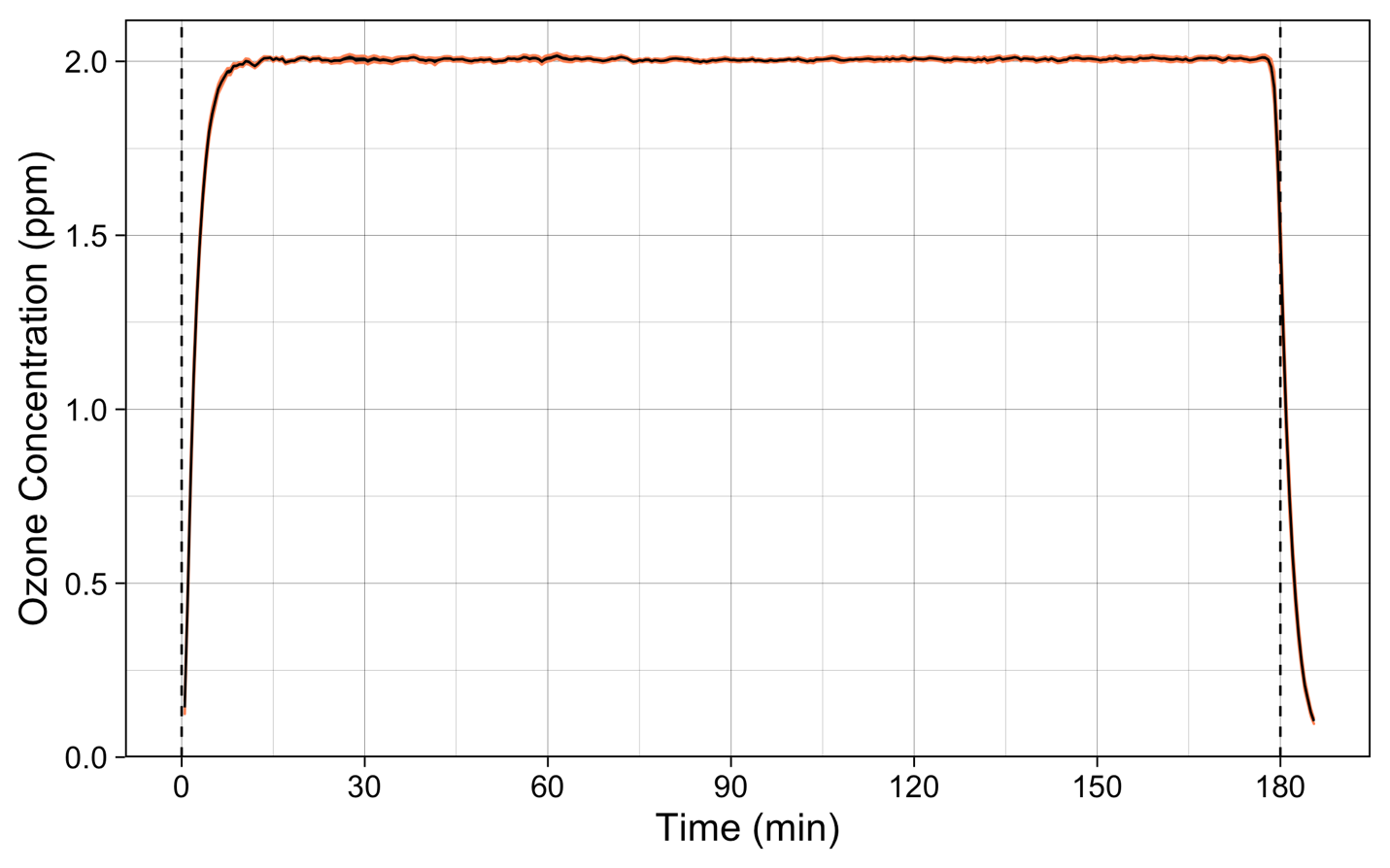

**Supplemental Figure S2. Two ppm ozone (O_3_) was precisely delivered across all exposures.** Mean (black line) and standard error (red shading) for synchronized exposure data across all 17 batches.

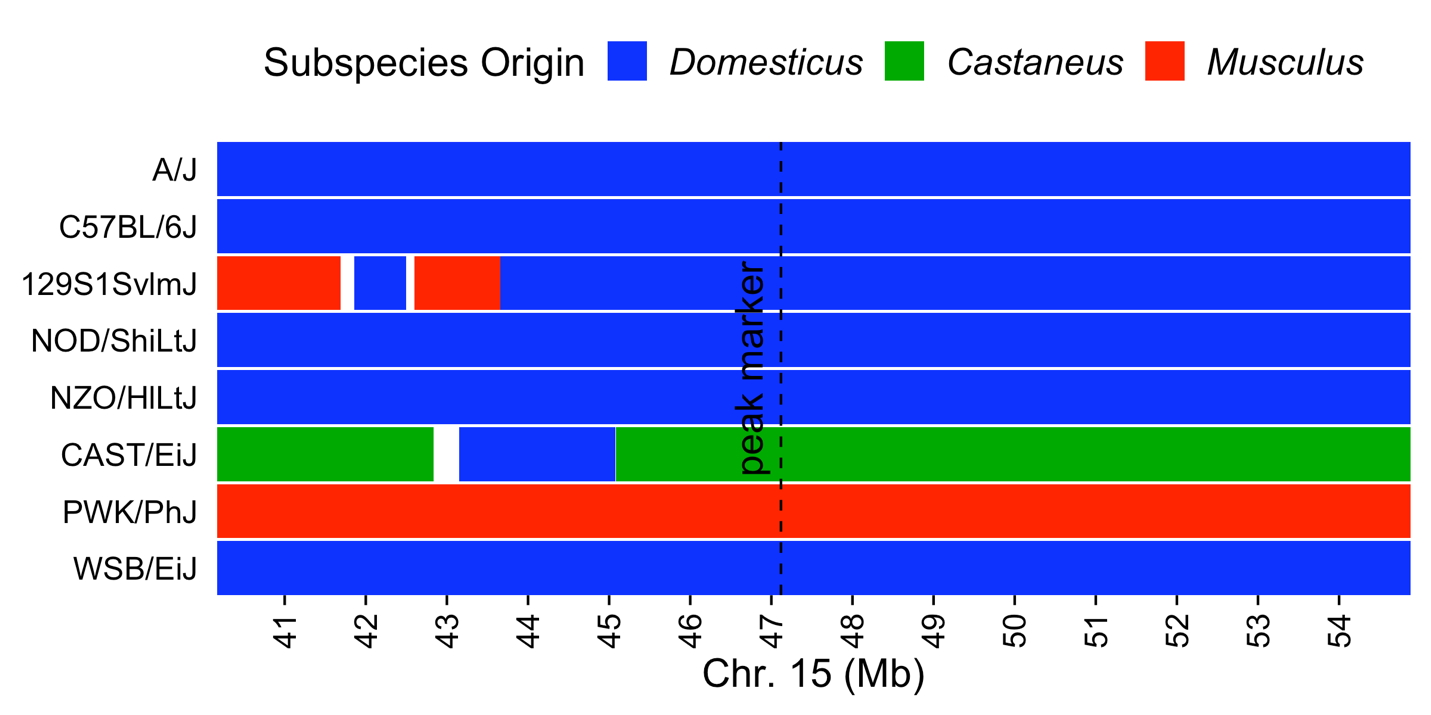

**Supplemental Figure S3. A ~2 Mb region of Mus musculus domesticus intersubspecific introgression is present in the CAST/EiJ genome near the peak marker within the Chromosome (Chr.) 15 QTL.** Subspecies origin for each of the 8 CC founder haplotypes within the span of the Chr. 15 QTL was visualized using the Mouse Phylogeny Viewer, and color-coded: Mus musculus domesticus is blue, M. m. castaneus is green, and M. m. musculus is red. White regions are haplotypes of undetermined origin.

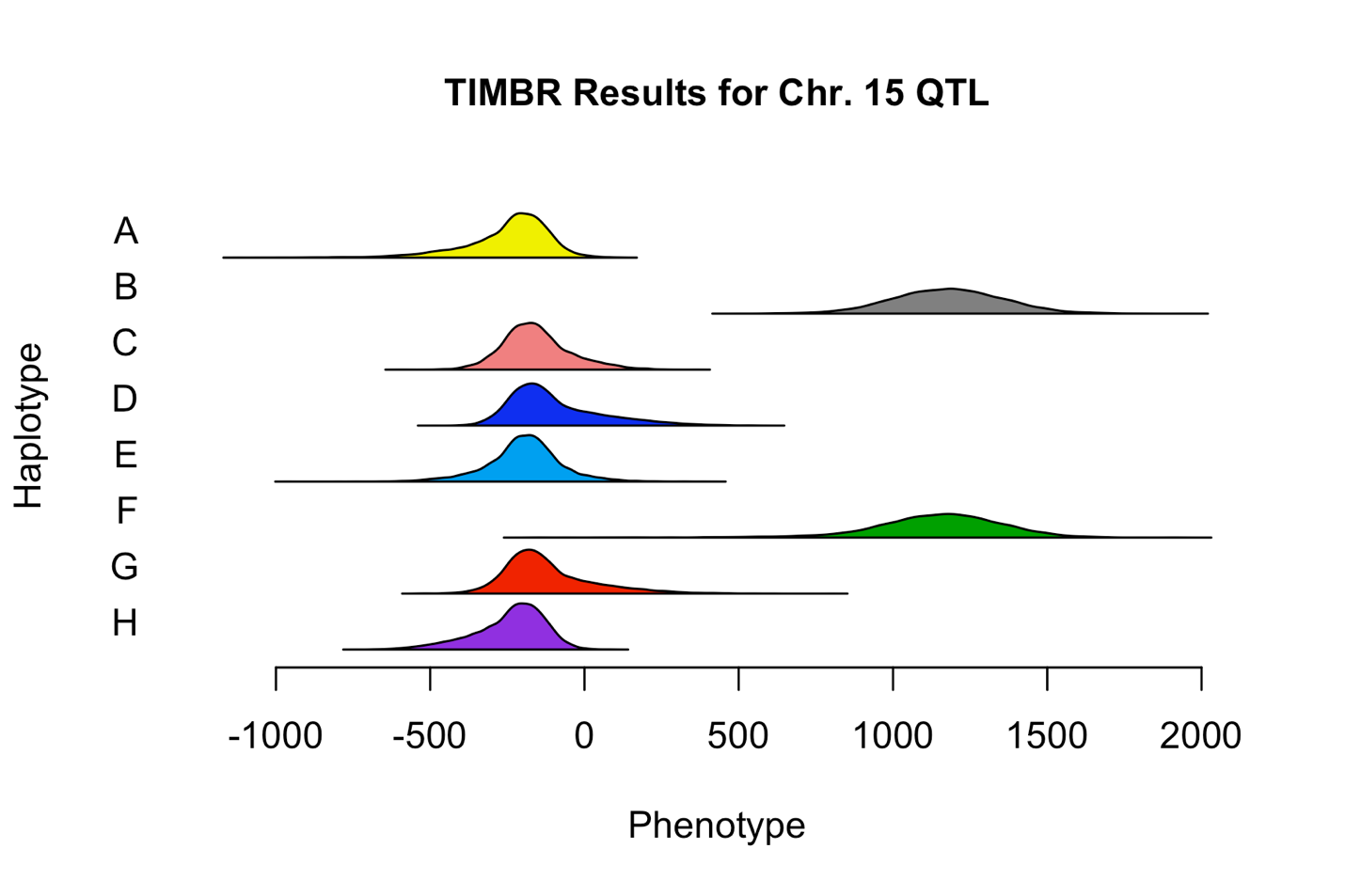

**Supplemental Figure S4. Posterior distribution of the haplotype effects estimated using TIMBR allelic series inference framework confirm grouping of C57BL/6J and CAST/EiJ.** A: A/J, B: C57BL/6J, C: 129S1SvImJ, D: NOD/ShiLtJ, E: NZO/ H1LtJ, F: CAST/EiJ, G: PWK/PhJ, H: WSB/EiJ.

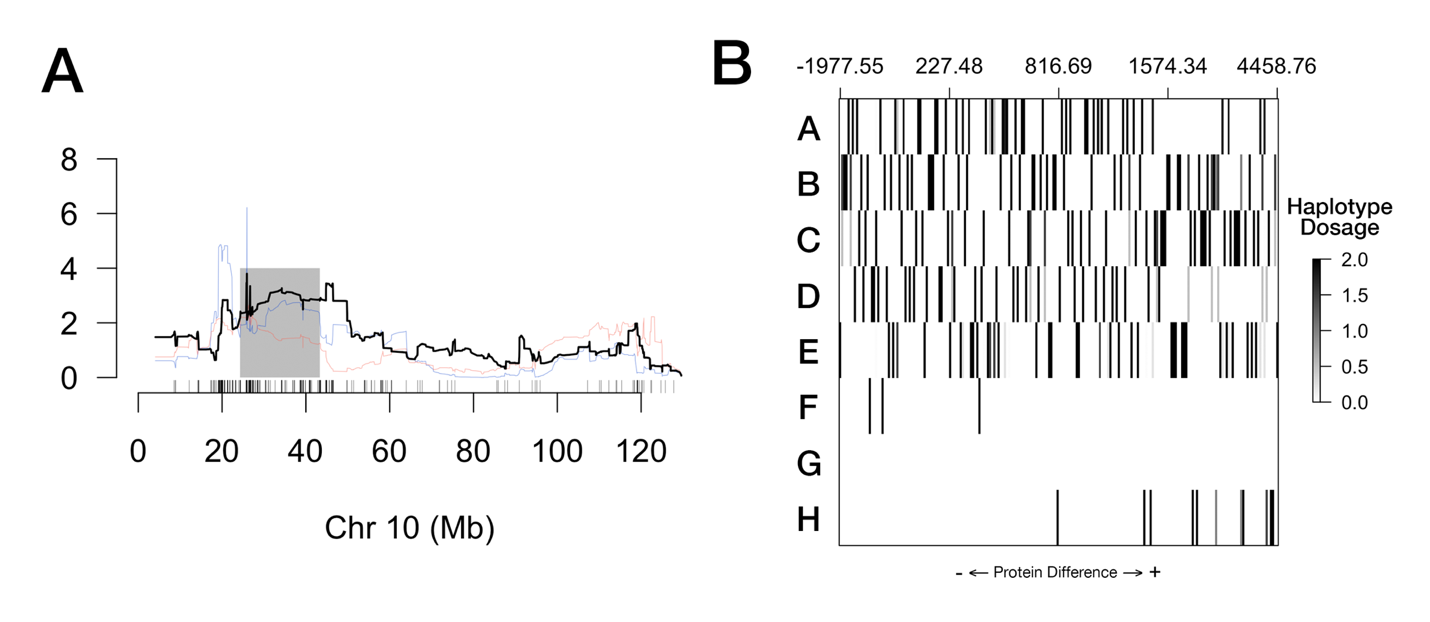
 **Supplemental Figure S5. A locus on Chromosome (Chr.) 10 is associated with total protein response, surpassing the 90% significance threshold.** (A) The locus *Oipq2* spans ~19 Mb on Chr. 10 (80% CI: 24.58-43.61 Mb), indicated by gray shading. The black line indicates genome scan resulting from QTL mapping, while the faint blue and red lines represent individual artificial scans resulting from the permutation procedure. (B) Strain matches with WSB/EiJ at the peak marker are associated with high protein response. A: A/J, B: C57BL/6J, C: 129S1SvImJ, D: NOD/ShiLtJ, E: NZO/ H1LtJ, F: CAST/EiJ, G: PWK/PhJ, H: WSB/EiJ.
